# Brain-Score: Which Artificial Neural Network for Object Recognition is most Brain-Like?

**DOI:** 10.1101/407007

**Authors:** Martin Schrimpf, Jonas Kubilius, Ha Hong, Najib J. Majaj, Rishi Rajalingham, Elias B. Issa, Kohitij Kar, Pouya Bashivan, Jonathan Prescott-Roy, Franziska Geiger, Kailyn Schmidt, Daniel L. K. Yamins, James J. DiCarlo

**Affiliations:** Department of Brain and Cognitive Sciences, MIT, Cambridge, MA 02139; Center for Brains, Minds and Machines, MIT, Cambridge, MA 02139; McGovern Institute for Brain Research, MIT, Cambridge, MA 02139; Brain and Cognition, KU Leuven, Leuven, Belgium; Bay Labs Inc., San Francisco, CA 94102; Center for Neural Science, New York University, New York, NY 10003; Department of Neuroscience, Zuckerman Mind Brain Behavior Institute, Columbia University, New York, NY 10027; Department of Psychology, Stanford University, Stanford, CA 94305; Department of Computer Science, Stanford University, Stanford, CA 94305

**Author notes:** Correspondence: * (M.S.), (J.K), (J.J.D.). Equal contribution (Martin Schrimpf, Jonas Kubilius).

**Keywords:** computational neuroscience, object recognition, deep neural networks

## Abstract

The internal representations of early deep artificial neural networks (ANNs) were found to be remarkably similar to the internal neural representations measured experimentally in the primate brain. Here we ask, as deep ANNs have continued to evolve, are they becoming more or less brain-like? ANNs that are most functionally similar to the brain will contain mechanisms that are most like those used by the brain. We therefore developed *Brain-Score* – a composite of multiple neural and behavioral benchmarks that score any ANN on how similar it is to the brain’s mechanisms for core object recognition – and we deployed it to evaluate a wide range of state-of-the-art deep ANNs. Using this scoring system, we here report that: (1) DenseNet-169, CORnet-S and ResNet-101 are the most brain-like ANNs. (2) There remains considerable variability in neural and behavioral responses that is not predicted by any ANN, suggesting that no ANN model has yet captured all the relevant mechanisms. (3) Extending prior work, we found that gains in ANN ImageNet performance led to gains on Brain-Score. However, correlation weakened at ≥ 70% top-1 ImageNet performance, suggesting that additional guidance from neuroscience is needed to make further advances in capturing brain mechanisms. (4) We uncovered smaller (i.e. less complex) ANNs that are more brain-like than many of the best-performing ImageNet models, which suggests the opportunity to simplify ANNs to better understand the ventral stream. The scoring system used here is far from complete. However, we propose that evaluating and tracking model-benchmark correspondences through a Brain-Score that is regularly updated with new brain data is an exciting opportunity: experimental benchmarks can be used to guide machine network evolution, and machine networks are mechanistic hypotheses of the brain’s network and thus drive next experiments. To facilitate both of these, we release Brain-Score.org: a platform that hosts the neural and behavioral benchmarks, where ANNs for visual processing can be submitted to receive a Brain-Score and their rank relative to other models, and where new experimental data can be naturally incorporated.

## Introduction

Deep convolutional artificial neural networks (ANNs) (LeCun et al., 2015) were derived in part from findings in visual neuroscience (see Yamins and DiCarlo (2016) for review) and are now the leading models in machine vision and other areas of AI. Soon after Krizhevsky et al. (2012)’s initial results, it was found that by evolving deep ANNs to achieve gains in performance (through either architecture search or weight training), those ANNs developed internal feature representations that are remarkably similar to neural representations recorded in mid and high levels of the non-human primate ventral visual processing stream (Yamins et al., 2013, 2014; Khaligh-Razavi and Kriegeskorte, 2014) (see Yamins and DiCarlo (2016) for review). More recent work has extended this same “performance-driven” ANN approach to the human visual system (Güçlü and van Gerven, 2015; Cichy et al., 2016; Kubilius et al., 2016), to lower levels of visual processing (Cadena et al., 2017), to auditory processing (Kell et al., 2018), and to the rodent tactile system (Zhuang et al., 2017).

The models from this early work outlined above outperformed all other neuroscience models at the time and yielded reasonable scores on predicting response patterns from both single unit activity and fMRI. It was also suggested that ANNs with improved task performance were likely to be even better matches to the primate visual stream (Yamins et al. (2014); Yamins and DiCarlo (2016)). We thus ask, as model performance has increased from Alexnet’s 57.67% top-1 on ImageNet to up to 85.4% (Mahajan et al., 2018)^1^ today, are these even better models of the primate visual stream?

To answer this question, we here propose *Brain-Score* to evaluate any ANN on how brain-like it is – focusing on the parts of the brain that have been implicated in visual object recognition. Brain-Score is a composite benchmark consisting of neural and behavioral benchmarks, where each benchmark refers to the application of a metric to a particular dataset. Neural metrics assess the similarity of internally observable signals: image-evoked feature activations in ANNs and image-evoked recorded neural activations in different primate brain regions. Behavioral metrics assess the similarity of the “outputs” of ANNs and primates, such as predictions on match-to-sample tasks. For this study, we assembled a base set of neural and behavioral benchmarks: neural recordings from cortical areas V4 and IT in macaque monkeys and behavioral data from humans. We then evaluated dozens of state-of-the-art deep ANNs on these three brain benchmarks and the resulting Brain-Score. We note that while neuroscience models have in the past been influential as building blocks of today’s deep neural networks, we do not claim that Brain-Score will automatically yield better models for machine learning. This is a benchmark for the brain sciences, encouraging a quantified evaluation of models on neural and behavioral data. Further, while benchmarking is common in machine learning, most of neuroscience still lacks standardized datasets and tools to perform such benchmarks routinely, and there is little awareness of the benefits of a rigorous evaluation of proposed models of brain mechanisms. Brain-Score is our attempt to bridge this gap and provide the means to track progress in brain-like model development on a scale that is larger than individual laboratories.

Our main contributions are the following:

- We replicate prior work (Yamins et al., 2014) showing that ANNs that have higher ImageNet performance tend to be more functionally similar to the ventral visual stream, and we extend that work by demonstrating that many state-of-the-art deep ANNs simultaneously score well on all three of the brain benchmarks (V4, IT, and behavior).
- We report that correlation between ImageNet performance and neural data prediction is weak for recent models (i.e. those with ImageNet top-1 performance ≥ 70%), but variability between models appears nontrivial. In other words, some ANNs appear to predict neural responses better than others but not because they perform better on ImageNet.
- We show that an ANN’s ImageNet performance correlates robustly with behavioral metrics, meaning that the image-by-image patterns of behavior of high-performing ANNs mostly resemble and predict those of primates. Similar to predicting neural response data, there is significant variability among ANNs, and the ANNs with the highest ImageNet performance (NAS-Net and PNASNet) predict behavioral data considerably worse than older models such as ResNet-101.
- We identify DenseNet-169, CORnet-S (a new shallow recurrent network) and ResNet-101 as the current top three models of the mechanisms underlying primate object recognition (under our current set of benchmarks).
- To enable fast evaluations of neural networks on brain data, we release a platform, Brain-Score.org, that hosts the neural and behavioral data and accompanying metrics, where ANNs for visual processing can easily be submitted to receive a Brain-Score and their rank relative to other models. The Brain-Score and the ranking of models will be updated regularly as new experimental benchmarks are added and new ANNs become available. We also plan to include anatomical benchmarks in a future update of Brain-Score. This platform can easily be extended with new data, such as human fMRI recordings.

Stepping back, we suggest that evaluating and tracking the anatomical, neural, and behavioral correspondences through a Brain-Score that is regularly updated with new brain data is an exciting opportunity: brain measurements can be used to guide ANN evolution to emulate brain functions not yet captured by ANNs and to make ANNs that are more humanlike in their patterns of success and failures. For the brain and cognitive sciences, the ANN that best emulates the brain simultaneously becomes the current best understanding of how the brain actually works, and the driver of next experiments.

## Brain Benchmarks

In the following section we outline the benchmarks that models are measured against. A benchmark consists of a metric applied to a specific set of experimental data, which here can be either neural recordings or behavioral measurements.

### Neural

The purpose of neural metrics is to establish how well internal representations of a source system (e.g., a neural network model) match the internal representations in a target system (e.g., a primate). Unlike typical machine learning benchmarks, these metrics provide a principled way to prefer some models over others even if their outputs are identical. We outline here one common metric, Neural Predictivity, which is a form of a linear regression.

### Neural Predictivity: Image-Level Neural Consistency

Neural Predictivity is used to evaluate how well responses X to given images in a source system (e.g., a deep ANN) predict the responses in a target system (e.g., a single neuron’s response in visual area IT). As inputs, this metric requires two assemblies of the form stimuli × neuroid where neuroids can either be neural recordings or model activations. First, source neuroids are mapped to each target neuroid using a linear transformation:

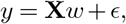

where *w* denotes linear regression weights and ϵ is the noise in the neural recordings. This mapping procedure is performed on multiple train-test splits across stimuli. In each run, the weights are fit to map from source neuroids to a target neuroid using training images, and then using these weights predicted responses *y*′ are obtained for the held-out images. We used the neuroids from V4 and IT separately to compute these fits. To obtain a neural predictivity score for each neuroid, we compare predicted responses *y*′ with the measured neuroid responses *y* by computing the Pearson correlation coefficient *r*:

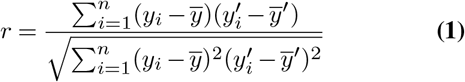

A median over all individual neuroid neural predictivity values (e.g., all measured target sites in a target brain region) is computed to obtain a predictivity score for that train-test split (median is used since responses are typically distributed non-normally). The final neural predictivity score for the target brain region is computed as the mean across all train-test splits.

We further estimate the internal consistency between neural responses by splitting neural responses in half across repeated presentations of the same image and computing Spearman-Brown-corrected Pearson correlation coefficient (Eq. 1) between the two splits across images for each neuroid.

In practice, we found that standard linear regression is comparably slow given a large dimensionality of the source system and not sufficiently robust. Thus, following Yamins et al. (2014), we use a partial least squares (PLS) regression with 25 components. We further optimized this procedure by first projecting source features into a lower-dimensional space using principal components analysis. The projection matrix is obtained for the features of a selection of ImageNet images, so that the projection is constant across train-test splits. This projection matrix is then used to transform source features. Results reported here were obtained by retaining 1000 principal components from the feature responses per layer to 1000 ImageNet validation images that captured the most variance of a source model.

### Neural Recordings

The neural dataset currently used in both neural benchmarks included in this version of Brain-Score is comprised of neural responses to 2,560 naturalistic stimuli in 88 V4 neurons and 168 IT neurons (cf. Fig. 1), collected by Majaj et al. (2015). The image set consists of 2,560 grayscale images in eight object categories (animals, boats, cars, chairs, faces, fruits, planes, tables). Each category contains eight unique objects (for instance, the “face” category has eight unique faces). The image set was generated by pasting a 3D object model on a naturalist background. In each image, the position, pose, and size of an object was randomly selected in order to create a challenging object recognition task both for primates and machines. A circular mask was applied to each image (see Majaj et al. (2015) for details on image generation).

**Figure 1:**
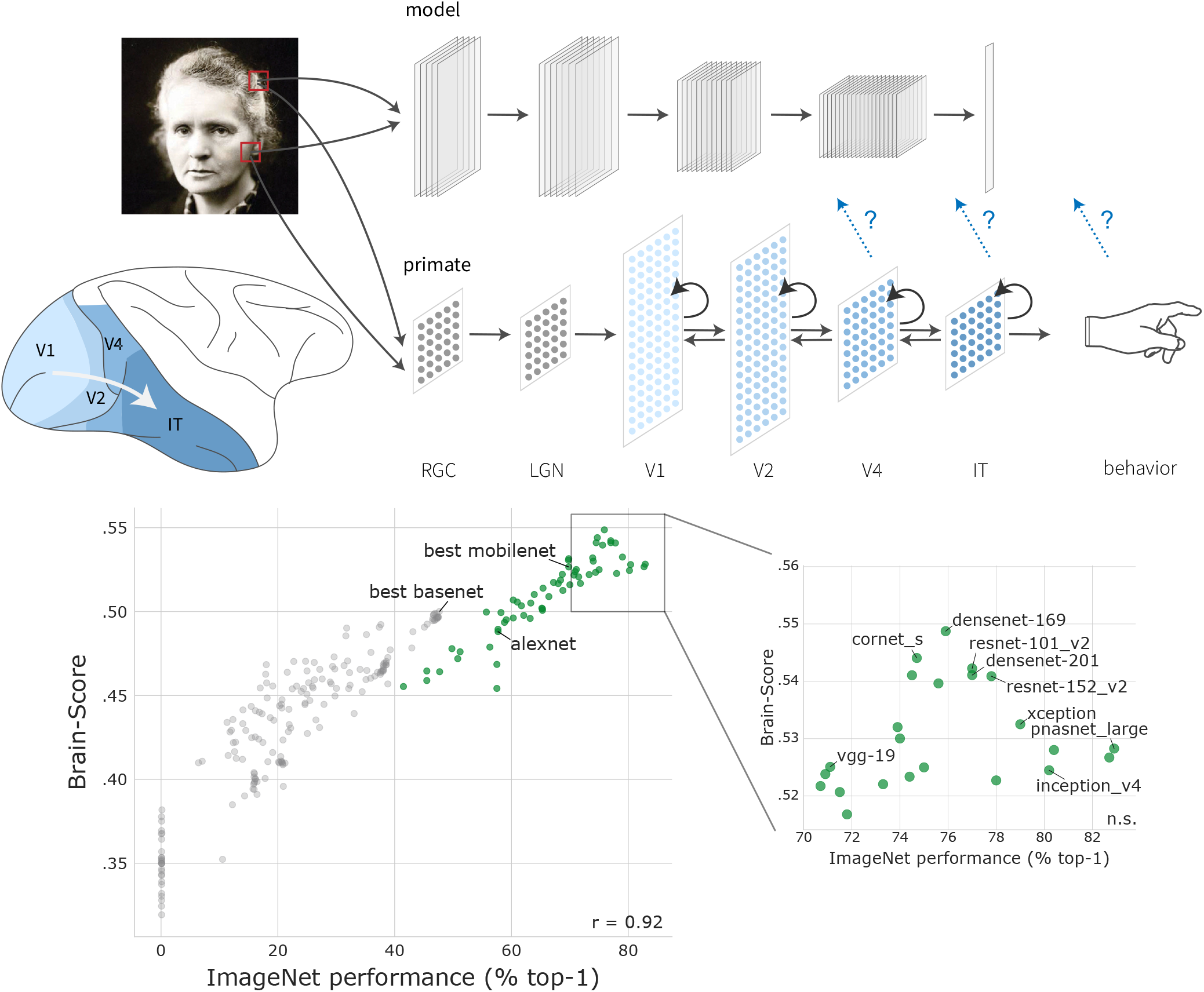
Overview of the Brain-Score. We compare neural networks using two classes of metrics: neural metrics compare the internal activations to regions of the macaque ventral stream, and behavioral metrics compare the similarity in outputs. Brain-Score is correlated with ImageNet performance for small, randomly combined models (gray dots) but becomes weak for current state-of-the-art models (green dots) at ≥ 70% top-1 performance.

Two macaque monkeys were implanted three arrays each, with one array placed in area V4 and the other two placed on the posterior-anterior axis of IT cortex. The monkeys passively observed a series of images (100 ms image duration with 100 ms of gap between each image) that each subtended approximately 8 deg visual angle. To obtain a stable estimate of the neural responses to each image, each each image was re-tested about 50 times (re-tests were randomly interleaved with other images). In the benchmarks used here, we used an average neural firing rate (normalized to a blank gray image response) in the window between 70 ms and 170 ms after image onset where the majority of object category-relevant information is contained (Majaj et al., 2015).

### Behavioral

The purpose of behavioral benchmarks it to compute the similarity between source (e.g., an ANN model) and target (e.g., human or monkey) behavioral responses in any given task. For core object recognition tasks, primates (both human and monkey) exhibit behavioral patterns that differ from ground truth labels. Thus, our primary benchmark here is a behavioral response pattern metric, not an overall accuracy metric, and higher scores are obtained by ANNs that produce and predict the primate patterns of successes and failures. One consequence of this is that ANNs that achieve 100% accuracy will not achieve a perfect behavioral similarity score.

Even within the visual behavioral domain of core object recognition, there are many possible behavioral metrics. We here use the metric of the image-by-image patterns of difficulty, broken down by the object choice alternatives (termed *I2n*), because recent work (Rajalingham et al., 2018) suggests that it has the most power to distinguish among alternative ANNs (assuming that sufficient amounts of behavioral data are available).

### I2n: Normalized Image-Level Behavioral Consistency

Source data (model features) for a total of *i* images are transformed first into a *i_b_* × *c* matrix of *c* object categories and *i_b_* images with behavioral data available using the following procedure. First, images where behavioral responses are not available (namely, *i* – *i_b_* images) are used to build a *c*-way logistic regression from source data to a *c*-value probability vector for each image, where each probability is the probability that a given object is in the image. This regression is then used to estimate probabilities for the held-out *i_b_* images. For each image, all normalized target-distractor pair probabilities are computed from the *c*-way probability vector. For instance, if an image contains a dog and the distractor is a bear, the target-distractor score is 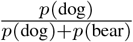.

In order to compare source and target data, we first transform these raw accuracies in the *i_b_* × *c* response matrix to a *d*′ measure for each cell in the *i_b_* × *c* matrix:

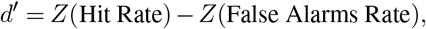

where *Z* is the estimated z-score of responses, Hit Rate is the accuracy of a given target-distractor pair while the False Alarms Rate corresponds to how often the observers incorrectly reported seeing that target object in images where another object was presented. For instance, if a given image contains a dog and distractor is a bear, the Hit Rate for the dog-bear pair for that image comes straight from the *i_b_* × *c* matrix, while in order to obtain the False Alarms Rate, all cells from that matrix that did not have dogs in the image but had a dog as a distractor are averaged, and 1 minus that value is used as a False Alarm Rate. All *d*′ above 5 were clipped. This transformation helps to remove bias in responses and also to diminish ceiling effects (since many primate accuracies were close to 1), but empirically observed benefits of *d*′ in this dataset are small; see Rajalingham et al. (2018) for a thorough explanation.

The resulting response matrix is further refined by subtracting the mean Hit Rate across trials of the same target-distractor pair (e.g., for dog-bear trials, their mean is subtracted from each trial). Such normalization exposes variance unique to each image and removes global trends that may be easier for models to capture. For instance, dog-bear trials on average could have been harder than dog-zebra trials. Without this normalization, a model might score very well by only capturing this tendency. After normalization, all responses are centered around zero, and thus capturing only global trends but not each image’s idiosyncrasies would be insufficient for a model to rank well.

After normalization, a Pearson correlation coefficient *r_st_* between source and target data is computed using Eq. 1. We further estimate noise ceiling, that is, how well an ideal model could perform given the noise in the measured behavioral responses, by dividing target data in half across trials, computing the normalized *d*′ *i_b_* × *c* matrices for each half, and computing the Pearson correlation coefficient *r_tt_* between the two halves. If source data is produced by a stochastic process, the same procedure can be carried out on the source data, resulting in the source’s reliability *r_ss_*.

The final behavioral predictivity score of each ANN is then computed by:

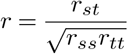

All models that we tested so far produced deterministic responses, thus *r_ss_* = 1 in our scoring.

### Primate behavioral data

The behavioral data used in the current round of benchmarks was obtained by Rajalingham et al. (2015) and Rajalingham et al. (2018). Here we focus on only the human behavioral data, but the human and nonhuman primate behavioral patterns are very similar to each other (Rajalingham et al., 2015, 2018).

The image set used in this data collection was generated in a similar way as the images for V4 and IT using 24 object categories. In total, the dataset contains 2,400 images (100 per object). For this benchmark, we used 240 (10 per object) of these images for which the most trials were obtained. 1,472 human observers responded to briefly presented images on Amazon Mechanical Turk. At each trial, an image was presented for 100 ms, followed by two response choices, one corresponding to the target object present in the image and the other being one of the remaining 23 objects (i.e., a distractor object). Participants responded by choosing which object was presented in the image. Thus, over three hundred thousand responses for each target-distractor pair were obtained from multiple participants, resulting in a 240 (images) × 24 (objects) response matrix when averaged across participants.

### Brain-Score

To evaluate how well a model is doing overall, we computed the global Brain-Score as a composite of neural V4 predictivity score, neural IT predictivity score, and behavioral I2n predictivity score (each of these scores was computed as described above). The Brain-Score presented here is the mean of the three scores. This approach does not normalize by different scales of the scores so it may be penalizing scores with low variance but it also does not make any assumptions about significant differences in the scores, which would be present in ranking.

## Candidate Models

For this round of evaluations, we sought to benchmark most commonly used neural network families: AlexNet (Krizhevsky et al., 2012), VGG (Simonyan and Zisserman, 2014), ResNet (He et al., 2015), Inception (Szegedy et al., 2015a, b, 2016), InceptionResNet (Szegedy et al., 2016), SqueezeNet (Iandola et al., 2016), DenseNet (Huang et al., 2017), MobileNet (Howard et al., 2017), and (P)NASNet (Zoph and Le, 2016; Liu et al., 2017). Most of pre-trained models were available in TensorFlow (Abadi et al., 2016), either via their Keras (Chollet et al., 2015) or Slim interface. For AlexNet, SqueezeNet, ResNet-18 and ResNet-34, we used their PyTorch implementation (Paszke et al., 2017).

To further map out the space of possible architectures and a baseline of neural, behavioral, and performance scores, we included an in-house-developed family of models with up to moderate ImageNet performance, termed *BaseNets*: lightweight AlexNet-like architectures with six convolutional layers and a single fully-connected layer, captured at various stages of training. Various hyperparameters were varied between BaseNets, such as the number of filter maps, nonlinearities, pooling, learning rate scheduling, and so on, and formed a basis for the CORnet family of models (Kubilius et al., 2018b).

We also tested CORnet-S, a new model that was developed with the goal of rivaling the best models on *Brain-Score* while being significantly shallower than competitors by leveraging bottleneck architecture and recurrence (Kubilius et al., 2018b). CORnet-S is composed of four recurrent areas with two to three convolutions each and a fully-connected layer at the end. For Neural Predictivities, we used activations at multiple internal layers of the networks. Layers were pre-selected by hand to include layers at multiple depths in each model and respecting the natural structuring (e.g., the outputs of a ResNet block were used, not the internal activations within the block). To keep the regression manageable, features were further downsampled with PCA to 1,000 dimensions. After testing every layer on both V4 and IT, we report the model’s score as the score of the best layer per region. Going forward, we are imagining more flexible methods for mapping model layers to brain regions, such as combining the activations of multiple layers. For CORnet-S, which already commits to a mapping to brain regions, we use the pre-defined mapping of the model. Behavioral scores were obtained using the final pre-readout layer of a network (i.e., the layer just before the last weight layer after which features are transformed into 1,000-dimensional outputs specific to the ImageNet task). In this case, features were not downsampled because typically dimensionality of the readout layer was sufficiently low to compute the scores quickly.

## Results

We examined a wide range of deep neural network trained on ImageNet and compared their internal representations with neural recordings in non-human visual cortical areas V4 and IT and with human behavioral measurements.

### Ranking of state-of-the-art networks

Table 1 summarizes the scores for each model on the range of brain benchmarks. The Brain-Score against ImageNet performance is shown in Fig. 1. The strongest model under our current set of benchmarks is DenseNet-169 with a Brain-Score of .549, closely followed by CORnet-S with a Brain-Score of .544 and ResNet-101 with a Brain-Score of .542. The current top-performing models on ImageNet from the machine learning community all stem from the DenseNet and ResNet families of models. DenseNet-169 and ResNet-101 are also among the highest-scoring models on the IT neural predictivity and the behavioral predictivity respectively with scores of .604 on IT (DenseNet-169, layer *conv5_block16_concat*) and .378 on behavior ResNet-101, layer *avg_pool*). VGG families win V4 with a score of .672 for VGG-19 (layer *block3_pool*).

**Table 1:** Brain-Scores and individual performances for state-of-the-art models

| Brain-Score | model | neural predictivity |  | behavioral predictivity | top-1 accuracy ImageNet |
| --- | --- | --- | --- | --- | --- |
|  |  | V4 | IT |  |  |
| **.549** | densenet-169 | .663 | **.606** | .378 | 75.90 |
| .544 | cornet_s | .650 | .600 | .382 | 74.70 |
| .542 | resnet-101_v2 | .653 | .585 | **.389** | 77.00 |
| .541 | densenet-201 | .655 | .601 | .368 | 77.00 |
| .541 | densenet-121 | .657 | .597 | .369 | 74.50 |
| .541 | resnet-152_v2 | .658 | .589 | .377 | 77.80 |
| .540 | resnet-50_v2 | .653 | .589 | .377 | 75.60 |
| .533 | xception | .671 | .565 | .361 | 79.00 |
| .532 | inception_v2 | .646 | .593 | .357 | 73.90 |
| .532 | inception_v1 | .649 | .583 | .362 | 69.80 |
| .531 | resnet-18 | .645 | .583 | .364 | 69.76 |
| .530 | nasnet_mobile | .650 | .598 | .342 | 74.00 |
| .528 | pnasnet_large | .644 | .590 | .351 | **82.90** |
| .528 | inception_resnet_v2 | .639 | .593 | .352 | 80.40 |
| .527 | nasnet_large | .650 | .591 | .339 | 82.70 |
| .527 | best mobilenet | .613 | .590 | .377 | 69.80 |
| .525 | vgg-19 | **.672** | .566 | .338 | 71.10 |
| .524 | inception_v4 | .628 | .575 | .371 | 80.20 |
| .523 | inception_v3 | .646 | .587 | .335 | 78.00 |
| .522 | resnet-34 | .629 | .559 | .378 | 73.30 |
| .521 | vgg-16 | .669 | .572 | .321 | 71.50 |
| .500 | best basenet | .652 | .592 | .256 | 47.64 |
| .488 | alexnet | .631 | .589 | .245 | 57.70 |
| .469 | squeezenet1_1 | .652 | .553 | .201 | 57.50 |
| .454 | squeezenet1_0 | .641 | .542 | .180 | 57.50 |

The best models from the BaseNet baseline family of models lag behind the winning models with a Brain-Score of .500 and a behavioral score of .256 but still perform reasonably well on V4 (.654) and IT (.592). Several observations for other model families are also worth noting: while ANNs from the Inception architectural family improved on ImageNet performance over subsequent versions, its Brain-Score decreased. Another natural cluster emerged with AlexNet and SqueezeNet at the bottom of the ranking: despite reasonable scores on V4 and IT neural predictivity, their behavioral scores are sub-par. Interestingly, models that score high on brain data are also not the ones ranking the highest on ImageNet performance, suggesting a potential disconnect between ImageNet performance and fidelity to brain mechanisms. For instance, despite its superior performance of 82.90% top-1 accuracy on ImageNet, PNASNet only ranks 13^th^ on the overall Brain-Score. Models with an ImageNet top-1 performance below 70% show a strong correlation with Brain-Score of .92 (*p* < 10^−14^) but above 70% ImageNet performance, there was no significant correlation (*p* >> .05, cf. Fig. 1). To investigate this potential disconnect further, we next analyzed the specific scores on neural and behavioral data.

### Scores on individual neural and behavioral benchmarks

Previous studies observed that models with higher classification performance tend to better predict neural data (Yamins et al., 2014). Here we extend that work by demonstrating that this performance-driven approach holds in a broad sense when evaluated on multiple deep neural networks in a wide range of ImageNet performance regimes, but fails to produce a network exactly matching the brain when reaching human performance levels (see Fig. 1). On individual scores, the correlation of ImageNet performance and Brain-Score varies substantially (Fig. 2). For instance, V4 single site responses are predicted best not only by VGG-19 (ImageNet top-1 performance 71.10%) but also Xception (79.00% top-1). Similarly, IT single site responses are predicted best by DenseNet-169 (.606; 75.90% top-1) but even BaseNets (.592; 47.64% top-1) and MobileNets (.590; 69.80% top-1) are very close to the same IT neural predictivity score. In contrast, the correlation between ImageNet performance and behavioral predictivity remains robust with AlexNet (57.50% top-1) or BaseNets performing substantially worse than the best models. However, the top-performing models on the behavioral score are not the state-of-the-art models on ImageNet: ResNet-101 ranks the highest on behavioral score (.389) but has 77.37% ImageNet top-1 performance, compared to PNASNet that achieves higher ImageNet performance (82.90% top-1) but a substantially lower behavioral score (.351). In fact, the correlation between ImageNet top-1 performance and behavioral score appears to be weakening, with models performing well on ImageNet exhibiting little correlation to behavioral scores, suggesting that better consistency with behavioral data might not be achieved by continuing the efforts to push ImageNet performance higher. Overall, despite the lack of clear trend at high ImageNet performance regimes, the performance-to-neural correlation is .68 (*p* < 10^−28^) in V4, .80 (*p* < 10^−47^) in IT, and performance correlates with behavior at .93 (*p* < 10^−91^).

**Figure 2:**
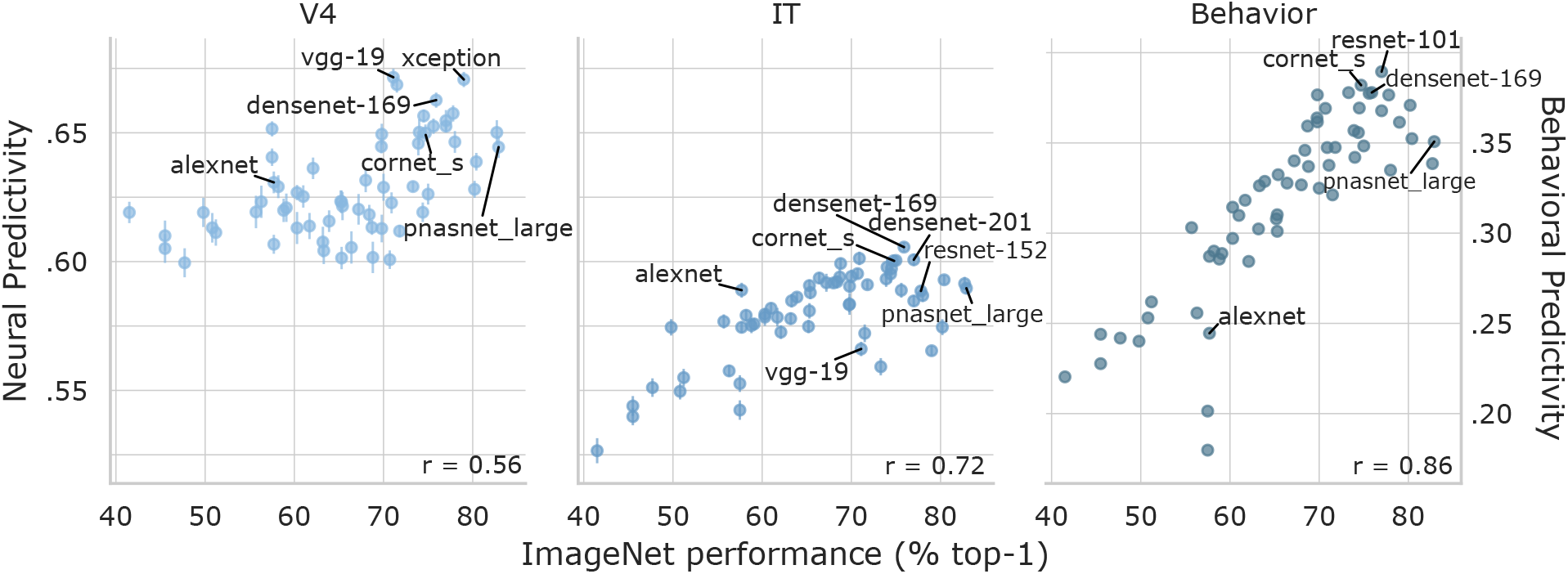
Predictivities of all models on neural and behavioral benchmarks. We evaluated regions V4 and IT using the *neural predictivity* as well as behavioral recordings using *I2n*. Current best models are: VGG-19on V4, DenseNet-169 on IT and ResNet-101 on behavior. Notably, DenseNet-169, CORnet-S and ResNet-101 are strong models on all three benchmarks. Noise ceilings are .892 for V4, .817 for IT and .497 for behavior. Error bars indicate s.e.m.

While all our current predictivity scores only summarize the average, it is clear that individual image-wise predictions are misaligned. In particular, out of the total 5,520 images, over half of the images (3,388, 61.38%) are already relatively well aligned between PNASNet (the best ImageNet model) and humans, as measured by no more than a 1*d*′ difference between human and model predictions. A substantial number of images (1,918, 34.75%) are easier for humans than models (Δ*d*′ > 1), meaning that further performance gains will simultaneously improve the behavioral score. However, on some images (214, 3.88%), the model outperforms humans (Δ*d*′ < −1) which might be desirable in a typical machine learning challenge but in fact hurts the model’s behavioral score as it tends to make the model more misaligned from humans.

We further analyzed the correlation of the neural scores to the behavioral score to determine the need for all individual benchmarks. We found that there was a moderate but not perfect correlation (.66 for behavior-to-V4 and .86 for behavior-to-IT) which provides justification for the full composite set of benchmarks outlined here. Put another way, this result confirms that even these first neural scores provide additional constraints on the mechanisms of the ventral stream beyond our high-resolution behavioral score.

Current models still fall short of reaching benchmark ceilings: The best ANN model V4 predictivity score is .663, which is below the internal consistency ceiling of these V4 data (.892). The best ANN model IT predictivity score is .604, which is below the internal consistency ceiling of these IT data (.817). And the best ANN model behavioral predictivity score is .378, which is below the the internal consistency ceiling of these behavioral data (.497).

## Discussion

We here present an initial framework for quantitatively comparing any artificial neural network to the brain’s neural network for visual processing. With even the relatively small number of brain benchmarks that we have included so far, the framework already reveals interesting patterns: It extends prior work showing that performance correlates with brain similarity, and our analysis of state-of-the-art networks yielded DenseNet-169, CORnet-S and ResNet-101 as the current best models of the primate visual stream. On the other hand, we also find a potential disconnect between ImageNet performance and Brain-Score: many of the best ImageNet models fall behind other models on Brain-Score, with the winning DenseNet-169 not being the best ImageNet model, and even small networks (“BaseNets”) with poor ImageNet performance achieving reasonable scores.

We do not believe that our initial set of chosen metrics is perfect, and we expect the metrics to evolve in several ways:

### By including more data of the same type used here

More neural sites collected with even the same set of images will provide more independent data samples, ensuring that models do not implicitly overfit a single set of benchmarks. Moreover, more data from more individuals will allow us to better estimate between-participant variability (i.e., the noise ceiling), establishing the upper bound of where models could possibly be (see below).

### By acquiring the same types of data using new images

Presently, our datasets use naturalistic images, generated by pasting objects on a random backgrounds. While these datasets are already extremely challenging, we will more stringently be able to test model ability to generalize beyond its training set by expanding our datasets to more classes of images (e.g., photographs, distorted images (Geirhos et al., 2018), artistic renderings (Kubilius et al., 2018a), images optimized for neural responses (Bashivan et al., 2018)).

### By acquiring the same types of data from other brain regions

The current benchmarks include V4, IT and behavioral readouts, but visual stimuli are first processed by the retina, LGN, V1 and V2 in the ventral stream. Including spiking neural data from these regions further constrains models in their early processing. Moreover, top-down modulation and control warrants recordings outside the ventral stream in regions such as PFC.

### By adding qualitatively new types of data

Our current set of neural responses consists of recordings from implanted electrode arrays, but in humans, fMRI recordings are much more common. Local Field Potential (LFP), ECoG, and EEG/MEG could also be valuable sources of data. Moreover, good models of the primate brain should not only predict neural and behavioral responses but should also match brain structure (anatomy) in terms of number of layers, their order, connectivity patterns, ratios of numbers of neurons in different areas, and so on. Finally, to scale this framework to a more holistic view of the brain, adding benchmarks for other tasks and domains outside of core object recognition is essential.

### By providing better experimental estimates of the ceilings of each component score

Note that it is still difficult to establish whether the ANN models are truly plateauing in their brain similarity – as implied in the results presented above – or if we are observing the limitations of our experimental datasets. For instance, neural ceilings only reflect the internal consistency of individuals neurons and, in that sense, are only an upper bound on the ceiling. That is, those neural responses are collected from individual monkeys, and it may be unreasonable to expect a single model to correctly predict every monkey’s neuron responses. A more reasonable ceiling might therefore need to reflect the consistency of an *average* monkey, leaving individual variabilities aside. However, in typical neuroscience experiments, recordings from only two monkeys are obtained, making it currently impossible to directly estimate these potentially lower ceilings.

Behavioral ceilings, on the other hand, might not be prone to such ceiling effects as they are already estimated using multiple humans responses (i.e. the “pooled” human data, see Rajalingham et al. (2015, 2018)). However, reaching consistency with the pooled human behavioral may not be the only way that one might want to use ANN models to inform brain science, as the across-subject variation is also an important aspect of the data that models should aim to inform on.

### By developing new ways to compute the similarity between models and data

Besides computing neural predictivity, there are multiple possible ways and particular parameter choices. Others have used for instance different versions of linear regression (Agrawal et al., 2014), RDMs (Khaligh-Razavi and Kriegeskorte, 2014; Cichy et al., 2016) or GLM (Cadena et al., 2017). We see neural predictivity as the current strongest form of comparing neural responses because it maps between the two systems and makes specific predictions on a spike-rate level. One could also use entirely new types of comparison, such as precise temporal dynamics of neural responses that are ignored here, even though they are likely to play an important role in brain function (Wang et al., 2018), or causal manipulations that may constrain models more strongly (Rajalingham and DiCarlo, 2018).

### By developing brain scores that are tuned separately for the non-human primate and the human

Our current set of benchmarks consist of recordings in macaques and behavioral measurements in humans and models are thus implicitly assumed to fit both of these primates. We do not believe that one ANN model should ultimately fit both species, so we imagine future versions of Brain-Score will treat them separately.

We caution that while Brain-Score reveals *that* one model is better than another, it does not yet reveal *why* that is the case. Due to current experimental constraints, we are not yet able to use Brain-Score to actually train a model. Both of these are key goals of our ongoing work.

To aid future efforts of aligning neural networks and the brain, we are building tools that allow researchers to quickly get a sense how their model scores against the available brain data on multiple dimensions, as well as compare against other models. Researchers can use our online platform Brain-Score.org to obtain all available brain data, submit new data and score their models on standardized benchmarks. The online platform provides an interface for submitting candidate models which are then automatically run on the current version of all benchmarks (code open-sourced at github.com/brain-score) and notify the submitting user about scores.

By providing this initial set of benchmarks we hope to ignite a discussion and further community-wide efforts around even better metrics, brain data and models. In this respect, our field is far closer to the beginning than the end, but it is important to get started and this is our version of such a start. We hope that Brain-Score will become a way of keeping track of computational models of the brain in terms of “how close we are” and quickly identifying the strongest model for a specific benchmark.

## Acknowledgments

This project has received funding from the European Union’s Horizon 2020 research and innovation programme under grant agreement No 705498 (J.K.), US National Eye Institute (R01-EY014970, J.J.D.), Office of Naval Research (MURI-114407, J.J.D), the Simons Foundation (SCGB [325500, 542965], J.J.D). This work was also supported in part by the Semiconductor Research Corporation (SRC) and DARPA.

## Author Contributions

M.S., J.P.R., F.G., and J.J.D. designed the platform. H.H., N.M., and K.S. collected neural data. E.B.I., K.K., R.R., and K.S. collected behavioral data. J.K., P.B., H.H., N.M., E.B.I., R.R., P.B., D.L.K.Y, and J.J.D developed metrics and benchmarks. M.S., J.K., H.H., J.P.R, and D.L.K.Y implemented metrics and benchmarks. M.S. and J.K. compared models. M.S, J.K., and J.J.D. wrote the paper.

## Footnotes

1 This model was unfortunately unavailable at the time of writing and we thus excluded it from the following analyses. The best Image Net model included here is PNASNet with 82.9% top-1 performance.

